# Daily Bowel Movements are Associated with Stronger Gut-Brain Phase-Amplitude Coupling and Better Cognitive Performance

**DOI:** 10.64898/2026.08.13.744397

**Authors:** Zeynep Ertürk, Klara Nielsen, Louise M.A. Jakobsen, Adam Duun Gottlieb, Hanne C. Bertram, Henrik M. Roager, Anke Ninija Karabanov

## Abstract

Gut–brain communication has emerged as a rapidly expanding field of research, with recent electrophysiological studies revealing rhythmic gut-brain coupling between gastric activity and brain oscillations in humans. Gut motility is a key determinant of gastrointestinal function, but it remains unclear whether individual differences in gut motility reflected by weekly bowel movements (e.g., defecation frequency) are associated with differences in gut-brain coupling. Here, we address this question by examining women with self-reported daily bowel movements (N = 38) and women with less frequent bowel movements (N = 38). We recorded simultaneous electroencephalography (EEG) and electrogastrography (EGG) at fasting state, performed cognitive assessments, and analysed faecal short-chain fatty acids (SCFAs) as markers of colonic fermentation. In a subset of participants, EEG-EGG coupling was assessed twice over an interval of at least eight weeks to assess test-retest reliability. In this group, EEG–EGG coupling showed moderate test–retest reliability (Intraclass Correlation Coefficient (ICC) = 0.50). When comparing the two groups of women, the phase-amplitude coupling (PAC) analysis between EEG and EGG signals revealed a significantly stronger gut-brain coupling in women with daily bowel movements compared to women with less frequent bowel movements (*p* = 0.03). We additionally found that women with daily bowel movements made less errors in the cognitive tasks and had higher levels of faecal SCFAs. A path analysis suggested that bowel movements significantly affect gut-brain phase-amplitude coupling through faecal SCFAs. However, neither faecal SCFAs nor phase-amplitude coupling significantly predicted cognitive performance, suggesting the existence of alternative pathways for the association between bowel movements and cognitive performance. Together, our findings suggest that the strength of gut–brain coupling is associated with bowel movements and cognitive performance, making EEG–EGG coupling a promising marker of human gut–brain interactions.

**Graphical Abstract:** 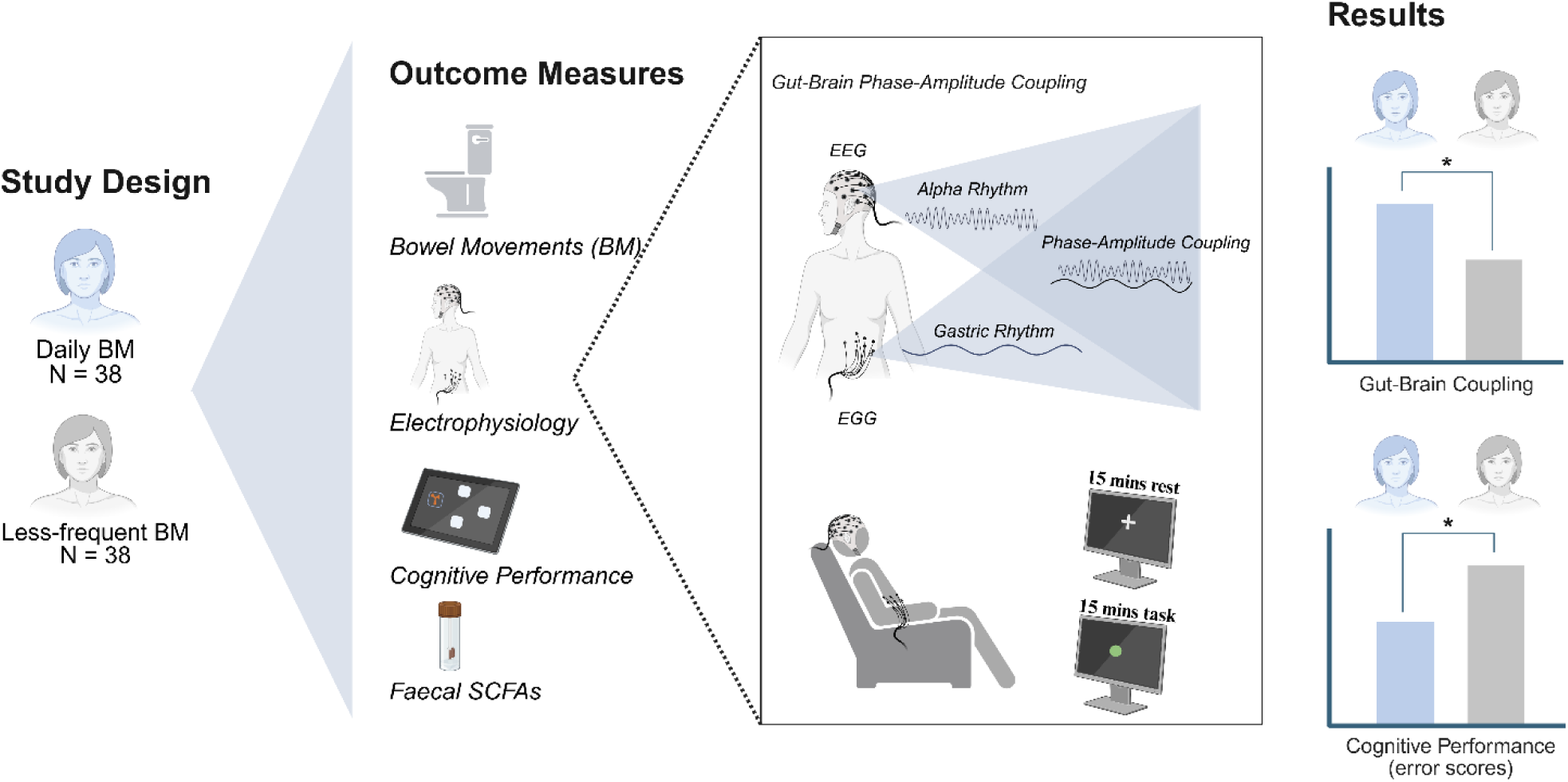

**Highlights:** Weekly bowel movement frequency is negatively associated with the global cognitive composite error score.

Gut-Brain Phase-Amplitude Coupling is positively associated with weekly bowel movement frequency.

Gut-Brain Phase-Amplitude Coupling is positively associated with levels of faecal short-chain fatty acids.

## Introduction

The gut-brain axis refers to the bidirectional communication between the gastrointestinal (GI) tract and the brain. Recent brain imaging studies have introduced a novel, non-invasive measure for gut-brain communication by showing significant coupling between gastric slow waves and brain activity in healthy humans. Furthermore, the strength of gut-brain coupling has been shown to link to cognitive and interoceptive states and mental health symptoms^1–4^ suggesting that gastrointestinal activity influences resting state brain activity networks^1,5^ as well as multisensory perception^6–8^, emotion^9^, working memory, and decision-making^10^. However, the physiological factors that shape gut-brain coupling remain poorly understood. One potential modulating factor is GI motility, as alterations in GI motility have been repeatedly associated with neuro-cognitive outcomes. Decreased bowel movement (BM) frequency has been associated with lower quality of life^11^ and worse cognitive function in cognitively healthy older adults^12^. In addition, an increasing body of work highlights a potential role of motility-related signals in brain dysfunction^13–15^, suggesting a connection between bowel movements, gut-brain coupling, and neuro-cognitive function.

At the physiological level, gastrointestinal motility is coordinated by interstitial cells of Cajal, which generate rhythmic electrical slow waves around 0.05 Hz in a healthy stomach^16^. These electrical slow waves contribute to the contractile activity of the GI tract and can be transmitted to the brain through afferent vagal pathways projecting to the nucleus tractus solitarii (NTS) and higher-order brain regions^17^. Alterations in gastric slow waves, including reductions in interstitial cells of Cajal density, have been observed in motility disorders such as slow-transit constipation^18,19^. It is, however, not known if such alterations also influence gut-brain coupling. In addition to electrophysiological motility signals, microbiome-derived chemical signalling may also modulate gut-brain coupling. Gut microbial metabolites such as short-chain fatty acids (SCFAs) influence enteric nervous system activity, bowel movements, and vagal tone^20,21^. Reduced levels of SCFA have been associated with bowel habits^22,23^, including slow-transit constipation^24^ and several neuropsychiatric disorders^25,26^, suggesting a potential biochemical pathway linking gut microbiome metabolic activity, GI motility, gut-brain coupling and cognition.

Electrogastrogram (EGG) is a non-invasive measure of the interstitial cells-generated slow waves and has long been used to characterise motility disorders^27,28^. Combining EGG with brain imaging has enabled the investigation of gut-brain coupling, defined as the synchronization between gastric slow waves and cortical activity. However, if natural variations in bowel movement frequency influence gut-brain coupling strength and if gut-brain coupling may mediate the association between low bowel movement frequency and cognition remains unexplored. Moreover, despite growing interest in gut-brain coupling as a marker of gut-brain communication, its consistency within individuals over time has been a source of concern, as some coupling metrics have been reported to have poor test-retest reliability^29^.

To address this knowledge gap, we first evaluated reproducibility of gut-brain phase-amplitude coupling strength across two experimental sessions to establish if phase-amplitude coupling can be used as a valid individual-level marker of gut-brain communication. Secondly, we investigated the connection between bowel movement frequency, cognition and phase-amplitude coupling by comparing cognitive function and phase-amplitude coupling between women with regular and women with infrequent bowel movements. Finally, we included a path analysis to explore potential biological mechanisms linking gut motility, cognitive function and gut-brain coupling, with faecal SCFA concentrations as a potential mediating factor.

## Methods Participants

In total, 40 women experiencing self-reported daily bowel movements (daily BM) (age: 45-65 years, BMI 16-38 kg/m2) and 56 women with self-reported less-frequent bowel movements (less-frequent BM) were included in the YourGutBrain project^30^ in Copenhagen, Denmark from April 2024 to November 2025 (ClinicalTrials.gov, NCT06311097). Exclusion criteria included pregnancy or lactation; a history of psychiatric, neurological, metabolic, or gastrointestinal disease; the use of antibiotics as well as medications known to affect bowel function or central nervous system activity (for the full list of Inclusion and exclusion criteria see **Appendix S1**). Participants were excluded because of withdrawal from the study (n = 17) or changes in the bowel movements frequency before the first experimental visit (n = 3). Consequently, a total of 38 women remained in the daily BM group, and 38 women in the less-frequent BM group. For the test-retest analysis in the less-frequent BM group, additional participants were excluded because of withdrawal before the re-test visit (n = 6) or poor EGG recording quality during the re-test visit (n = 1) resulting in a final sample set of 31 women in the test-retest group. The study was approved by the Municipal Ethical Committee of the Capital Region of Denmark (H-23071580) and participants received written and oral information about the study and provided written informed consent before participation in compliance with the principles of the Declaration of Helsinki. Participants received a voucher as compensation for their time.

## Study Design and Procedures

The current study presents baseline data from the YourGutBrain trial. The results of the intervention trial, investigating the effects of fermented dairy consumption on bowel movements and cognitive function, will be reported elsewhere. The trial includes two sets of baseline data: data of the daily-BM group that completed a single visit (V1), and data of the less-frequent BM group that completed four visits in a randomized crossover design, with a ≥ 4-week washout period between interventions. In the less-frequent BM group, V1 and V3 represent baseline pre-intervention visits, and V2 and V4 were post-intervention visits (**Figure 1A**). Here, we used V1 and V3 data to assess test–retest reliability of gut–brain coupling, and V1 data from both groups to investigate the links between bowel movement frequency, cognition, and gut–brain coupling. At each visit, all participants completed an identical test battery including a defecation diary, stool samples, cognitive assessments, questionnaires assessing physical and mental well-being, and electrophysiological recordings of gut–brain coupling. All measures are described in detail below.

**Figure 1.**
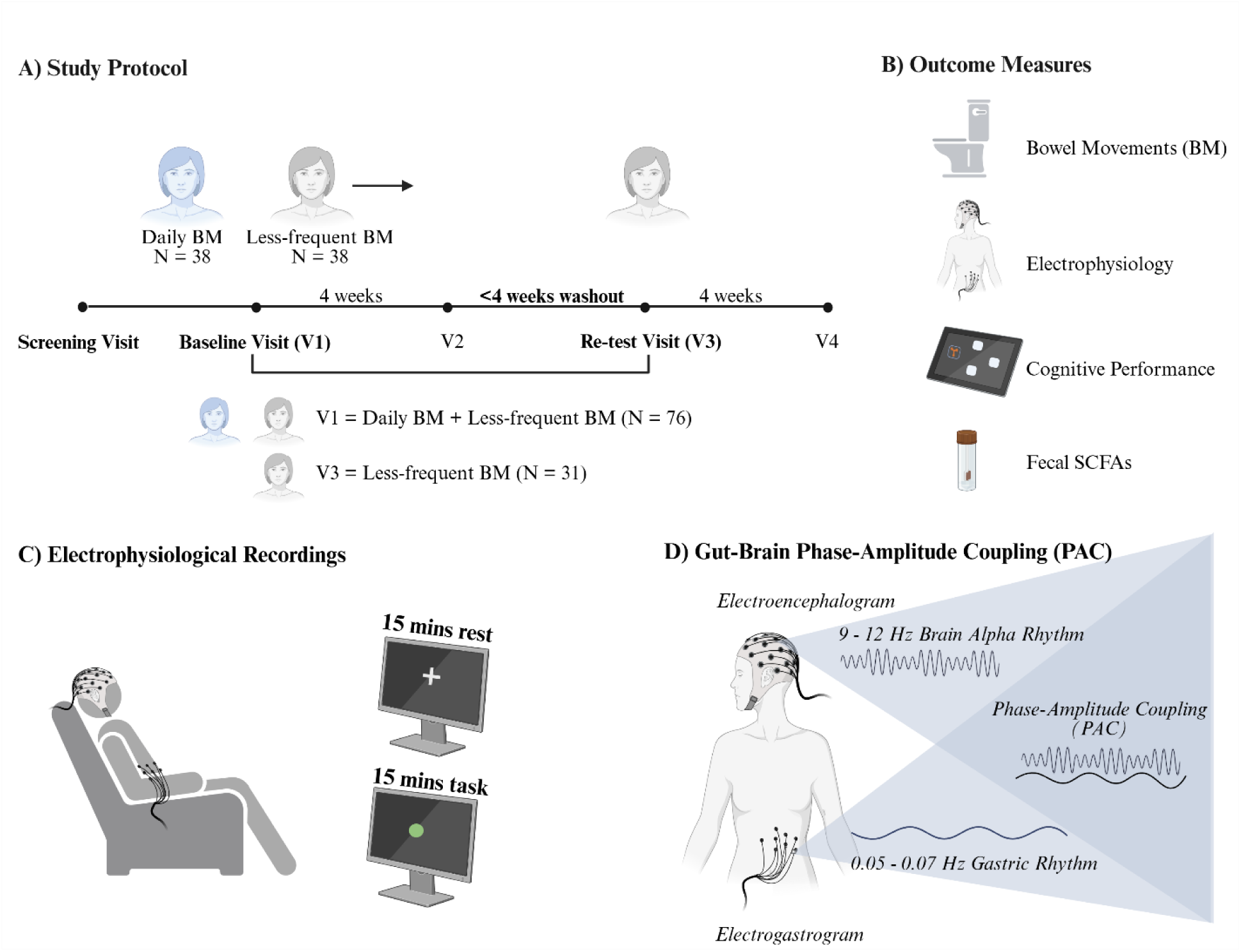
Overview of the stud design and assessments. (A) Study protocol illustrating the baseline comparison between the daily bowel movement (BM) group and the less-frequent BM group, as well as the test–retest reliability assessment between the baseline visit (V1) and the re-test visit (V3) in the less-frequent BM group. (B) Outcome measures collected at each study visit. (C) Simultaneous EEG and EGG recordings during both rest and task conditions. (D) Phase modulation of the gastric rhythms on the brain alpha rhythms is outlined. Participants were fasting during each visit, and all visits took place in the morning.

## Weekly bowel movements and cognitive assessments

*Bowel Movements:* Number of bowel movements was self-recorded using the Defecation Diary (**Appendix S2**) during the week before each visit. Participants wrote down the date and the time of defecations for one week, providing the number of weekly bowel movements. Bowel movement frequency was determined by counting the number of registered defecations for each participant during the recording period.

### Questionnaires

During the screening visit, participants’ socioeconomic status was assessed using self-reported education level, household income and employment status. Additionally, menopausal

symptoms were assessed using a questionnaire including somatic, psychological and urogenital items (**Appendix S3**).

At each experimental visit, the Pittsburgh Sleep Quality Index (PSQI)^31^ was used to assess subjective sleep latency, duration, efficiency and disturbances. Physical activity levels were evaluated using the International Physical Activity Questionnaire (IPAQ)^32^. Negative emotional states of depression, anxiety, and stress were assessed using DASS-42^33^, while health-related Quality of Life was assessed using the 36-Item Short Form Survey (SF-36)^34^. For sleep, a summed global PSQI score was generated following the original PSQI guidelines with higher scores indicating poorer sleep quality. For physical activity, the IPAQ was converted to metabolic equivalent scores (MET) following the standardized IPAQ scoring protocol^32^. For mental health, the DASS-42 Subscale scores were calculated as the sum of their respective Likert scale items following the DASS-42 scoring guidelines^33^, with higher scores indicating greater symptom severity. A total DASS score was computed as the sum of all subscales. For quality of life, SF-36 responses were evaluated on a 0-100 scale following the SF-36 guidelines^34^, with higher scores indicating better health status. Finally, the sum of the menopause questionnaire corresponding items was calculated as domain scores, with higher scores indicating higher symptom severity.

### Cognitive Tests

Motor function, memory, attention, and emotional bias were assessed using CANTAB (Cambridge Cognition, Cambridge, UK) using the Motor Screening, Paired Associates Learning, Rapid Visual Information Processing, and Emotional Bias Tasks. Each cognitive test was conducted on an iPad screen. Two participants were excluded from the cognitive score analyses: one due to feeling unwell during testing and the other due to additional task exposure resulting from a repeated study visit.

A composite cognitive score was calculated (composite-z) using the reaction time from the motor screening task and the error measures from each domain excluding the emotional bias task, representing the average global cognitive performance. This was done by log-transforming and standardizing values into z-scores within each measure across all participants and finally computing the composite score as the mean of the z-scores across the selected measures. As a known confounding factor in cognitive assessments^35^, socioeconomic status was adjusted for CANTAB analyses. Education and income were standardized and combined into a composite socioeconomic status score. Linear regression models were used to adjust for SES. CANTAB composite z-score is reported both before and after adjustment for socioeconomic status.

## Faecal short-chain fatty acids

Participants collected a faecal sample before each experimental visit. The samples were stored in airtight containers in a freezer at home until the day of the experimental visit. Cooling bags and elements were used for transportation to the study site. The samples were stored at -20 °C freezer until further processing. The frozen faecal samples were thawed at 4 °C overnight, diluted 1:1 (w/v) with pure Milli-Q water, homogenized and aliquoted in 1 mL cryotubes and stored at -70 °C.

The samples were subsequently analysed by proton nuclear magnetic resonance (^1^H-NMR) spectroscopy to quantify the individual faecal metabolite profiles as previously reported^36^.In brief, faecal samples were thawed and 400 µL faecal slurry was mixed 1:1 (v/v) with phosphate buffer (75 mM, 0.6 mM 3-(trimethylsilyl)-propionic-2,2,3,3-d4 acid (TSP), 20 % D_2_O) and 200 µL Milli-Q water on ThermoMixer (Eppendorf) for 2 minutes. Samples were centrifuged for 20 minutes (14.000g at 4 °C) and 600 µL supernatant was transferred to 5 mm NMR tubes. ^1^H NMR spectroscopy was conducted at 300 K on 14 T Bruker Avance III spectrometer (Bruker BioSpin, Rheinstetten, Germany) equipped with a 5 mm probe, automated tuning and matching accessory (ATMATM), BCU-I for the regulation of temperature, and SampleJet robot cooling system set to 5°C as a sample changer. ^1^H NMR spectra were acquired using NOESY pre-saturation pulse sequence (Bruker 1D noesygppr1D), 64 K data points, spectral width of 20 ppm, an acquisition time of 2.75 s, a relaxation delay of 4 s, 64 scans, and a fixed receiver gain. Quantification of short-chain fatty acids were achieved using Chenomx software (version 10.0 Professional, Chenomx Inc. Edmonton, Canada) to integrate and quantify the metabolite peaks relative to the TSP standard. Concentrations of the most common SCFAs (acetate, propionate, butyrate) were determined for each participant and used for all subsequent analyses.

## Electrophysiological Recordings

Participants arrived fasting (fasted from 9 pm previous night) in the morning to each visit to avoid digestion-related artifacts of muscle and movement artifacts on the abdominal region. Simultaneous 64-electrode EEG BioSemi ActiveTwo was used to acquire brain activity (Biosemi Inc., Amsterdam, The Netherlands) and 8 BioSemi external electrodes were used to acquire gastric activity. The external electrodes were placed on the abdominal region using an unipolar EGG montage^37^. Scalp and abdominal skin were cleaned and prepared before electrode position to ensure low impedance. Recordings for each visit included a continuous 15-minute recording of eyes-open resting condition to capture the spontaneous gut-brain coupling during rest. Participants were instructed to avoid structural mental processes, including counting or mentally repeating a text. The resting state recording was followed by a continuous 15-minute task recording where participants performed a response inhibition task (“Go-No-Go” task) during the electrophysiology recording. This was added to capture changes in gut-brain coupling during a task that required an external focus.

### EEG Preprocessing

All electrophysiological data were analysed using MATLAB (Mathworks, USA, Version R2023b), EEGLAB^38^ (version 2025.1.0) and fieldTrip^39^ (version 20231220). EEG data was filtered for the alpha band (8-12 Hz). The continuous recordings were visually inspected for bad channels. Independent component analysis (ICA) was computed using runica function on EEG channels. Components were visually inspected and muscle, eye, cardiac components were removed. Alpha activity was assessed across peak alpha region and peak alpha frequency (PAF) between groups using Welch’s independent samples t-test. Statistical significance was set at *p* < 0.05.

### EGG Preprocessing

EGG preprocessing and quality control steps followed the recent recommendations^37^. Each participant’s EGG peak within the normogastria range was identified on a power spectrum. Selected channel was bandpass filtered using a FIR band centred on the peak frequency with bandwidth -+0.015 Hz. Phase of the signal was extracted using Hilbert Transform. Visual inspection of each signal confirmed a plausible dominant rhythm, filtering behaved as expected, and no gross artifacts remained. Standard deviation of cycle duration was used as a stability indicator. The index of individual peak information for each participant was saved for the gut-brain phase-amplitude coupling analysis. Group differences in maximum EGG power within the monogastric range and selected peak electrode count were assessed using Welch’s independent-samples t-test. Statistical significance was set at *p* < 0.05.

### PAC Analysis

To quantify gut-brain coupling, we adopted Phase-amplitude coupling analysis^40^ to quantify the modulation effect of the gastric phase on the amplitude of the cortical alpha rhythm using the Modulation Index (MI) as the coupling measure. The analytical phase of EGG and amplitude envelope of EEG alpha-band were obtained using the Hilbert transform. The gastric phase cycles were divided into 18 equally spaced phase bins (-pi pi)^41^. The mean alpha amplitude was computed for each bin, creating a phase amplitude distribution. This distribution was normalized to a probability distribution, and Shannon entropy was used to calculate MI reflecting the deviation of the alpha amplitude distribution across the gastric phase bins from a uniform distribution. MI was computed for each EEG channel, returning one MI value between selected individual gastric signal and each 64 EEG signal. Grand-average MI topographies were generated by averaging MI values across the group.

### Significance Analysis of Phase Amplitude Coupling

The statistical procedure for identifying significant phase amplitude coupling at the group level followed previous studies^42^. Cross-person surrogate MIs were computed by pairing the individual EEG signals with gastric phase signals from all other participants as surrogates (N = 76). The median surrogate MI was calculated for each electrode and subtracted from the empirical MI to obtain ΔMI. Group-level significance of ΔMI was evaluated using a two-sided, cluster-based permutation test with 10,000 permutations. Clusters were defined based on a cluster-forming threshold (*p* < 0.005) and six nearest neighbour electrodes in two-dimensional sensor space. MI Cluster significance was determined using family-wise error correction (*p* < 0.05). Electrodes belonging to the significant contiguous MI cluster identified across all participants’ V1 visits were defined as Region of Interest (ROI), and the mean empirical MI across these electrodes was calculated for each participant for subsequent analysis.

## Statistical Analyses

All statistical analyses were performed in R^43^ (version 4.5.2). Data are presented as mean and standard deviation (SD). Normality of continuous variables was assessed using Shapiro-Wilk test. Statistical significance was defined as *p* < 0.05.

### Outlier Handling and Missing Data Imputation

In faecal SCFAs, potential outliers were identified through visual inspection of boxplots and subsequently evaluated by reviewing the original NMR spectra to identify whether they represent biologically plausible values. One sample was identified as an outlier and excluded from the primary analysis. The median metabolite concentrations were consistent with the means calculated after exclusion, indicating that the outlier did not substantially affect the results. For all other measures, no additional outliers were identified or removed. Education and income data were missing for some participants (N=7) and were handled using multiple imputations with predictive mean matching (20 imputations). A composite socioeconomic status score was then calculated within each imputed dataset and included as a covariate in the adjusted analyses of cognitive performance. None of the other measures required missing data imputation.

### MI Test-retest reliability

Test-retest reliability of the MI was evaluated at both V1 and V3 in the less-frequent BM group. We first evaluated the consistency of the selected peak EGG electrode within the normogastria range and the EEG alpha power by comparing measurements obtained at the baseline and test-retest visits. Changes in EEG peak alpha power were assessed using paired-sample *t*-test. The stability of the selected EGG peak channel was evaluated by calculating the percentage of participants for whom the same channel was identified at both visits. Test-retest reliability of EEG peak alpha power and peak EGG gastric frequency was quantified using intraclass correlation coefficients (ICCs). Changes in MI across all the significant electrodes were assessed using paired Wilcoxon signed-rank test and the test-retest reliability of MI was assessed through ICC using the MI measurements across all significant electrodes. For all the further group analysis containing MI, the ROI included the largest contiguous cluster, which resulted in exclusion of a single electrode.

### Statistical Group Comparisons

Bowel movement frequency, MI, Global PSQI scores, DASS-42, IPAQ, SF-36 scores, and CANTAB composite z-score were compared between groups using Wilcoxon rank-sum tests.

### Correlation Analyses

Spearman correlations were performed to examine relationships between bowel movements, cognitive performance, phase-amplitude coupling, and faecal short chain fatty acids. Correlation coefficients and corresponding p-values were calculated. To account for multiple testing, FDR was controlled with Benjamini Hochberg^44^ procedure for the multiple comparisons of SCFA abundance, and for the Spearman correlation analyses between different SCFA and MI.

### Mediation Analyses

Mediation analysis was performed using path analysis implemented in the R package lavaan^45,46^. We examined whether the association between bowel movements and cognitive performance was mediated by SCFA and MI. Total SCFA was computed as the sum of acetate, propionate, and butyrate. Direct, indirect and total effects were estimated simultaneously within a single structural equation model. Standard errors and 95% confidence intervals were obtained using nonparametric bootstrapping with 5000 iterations.

## Result

### Moderate test re-test reliability for modulation index

The ICC analysis revealed a moderate test-retest reliability of MI in the significant cluster across visits (ICC (3,1) = 0.50, 95% CI [0.18, 0.72]) (**Figure 2A**). The MI across the significant cluster was higher in the retest visit (*p* = 0.05) (**Figure 2B**). The ICC analysis further revealed good test-retest reliability of the alpha power (ICC (3,1) = 0.89, 95% CI [0.78, 0.94]) (**Figure S1**) and moderate reliability of peak gastric frequency (ICC (3,1) = 0.55, 95% CI [0.25, 0.76]) (**Figure S2**). The selected EGG electrode showed 51.6% exact agreement between the visits (**Figure S3**). We observed an overall within-participant stability of the EEG alpha power (**Figure 2C**) and selected peak EGG channel across visits (**Figure 2D**). At both visits, there was a similar spatial distribution of the peak gastric electrode selected for the MI analysis, reflecting the within-participant regional stability of the peak gastric activity. Although there were no significant BM differences within participants across visits (*p* = 0.09), BM was observed to increase with an average of 0.55 units (SD = 1.77) from visit 1 (baseline) to visit 3 in the 32 participants with few weekly BM who completed both visits.

**Figure 2.**
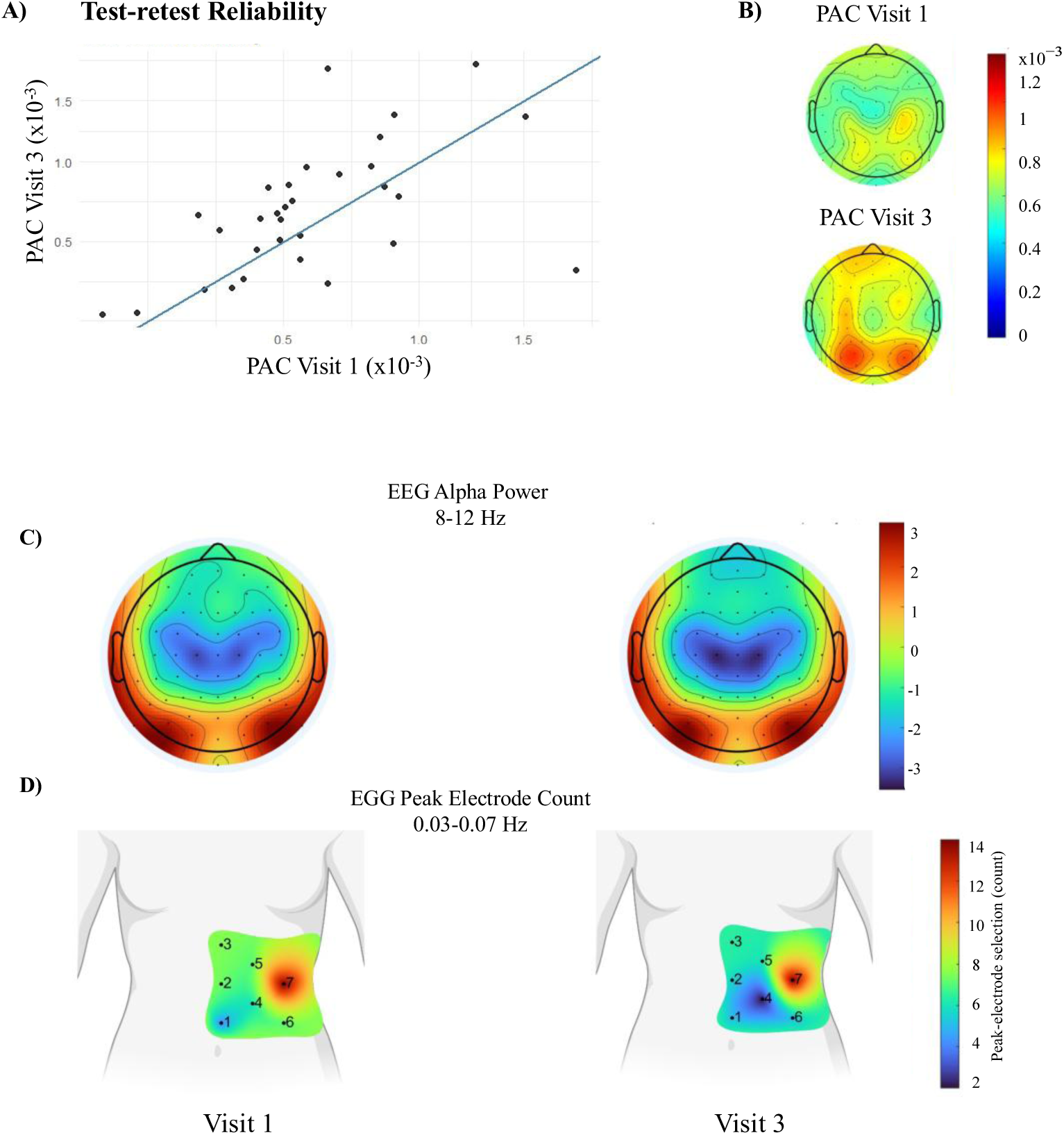
Modulation index test-retest reliability. (A) Scatter plot of modulation index (MI) values measured at baseline (V1; x-axis) and retest (V3; y-axis), illustrating test–retest reliability. (B) Topographical map showing the differences in MI between baseline (V1) and retest (V3). (C) Topographical distributions of mean EEG alpha power at baseline (V1) and retest (V3). (D) Topographical distributions of EGG peak electrode count at baseline (V1) and retest (V3).

### Groups with different BM frequency show similar baseline characteristics

The two groups of women differed as expected in bowel movement frequency (CI = [-7.00 -5.00], *p* < 0.001), while the groups were comparable with respect to age, BMI, physical activity, sleep quality, menopause symptoms, psychological symptoms, and quality of life (**Table 1**). Although emotional well-being scores were high and within the healthy range in both groups, a significant difference between groups was noted with less-frequent BM group having a higher rating of emotional well-being (CI = [-8 0], *p* = 0.04). Among the less-frequent BM group, 23 women were post-menopausal, six were pre-menopausal, and ten did not know. Among the daily BM group, 28 were post-menopausal, six were pre-menopausal, and three did not know. There was no significant difference in menopause symptoms between groups.

**Table 1.**
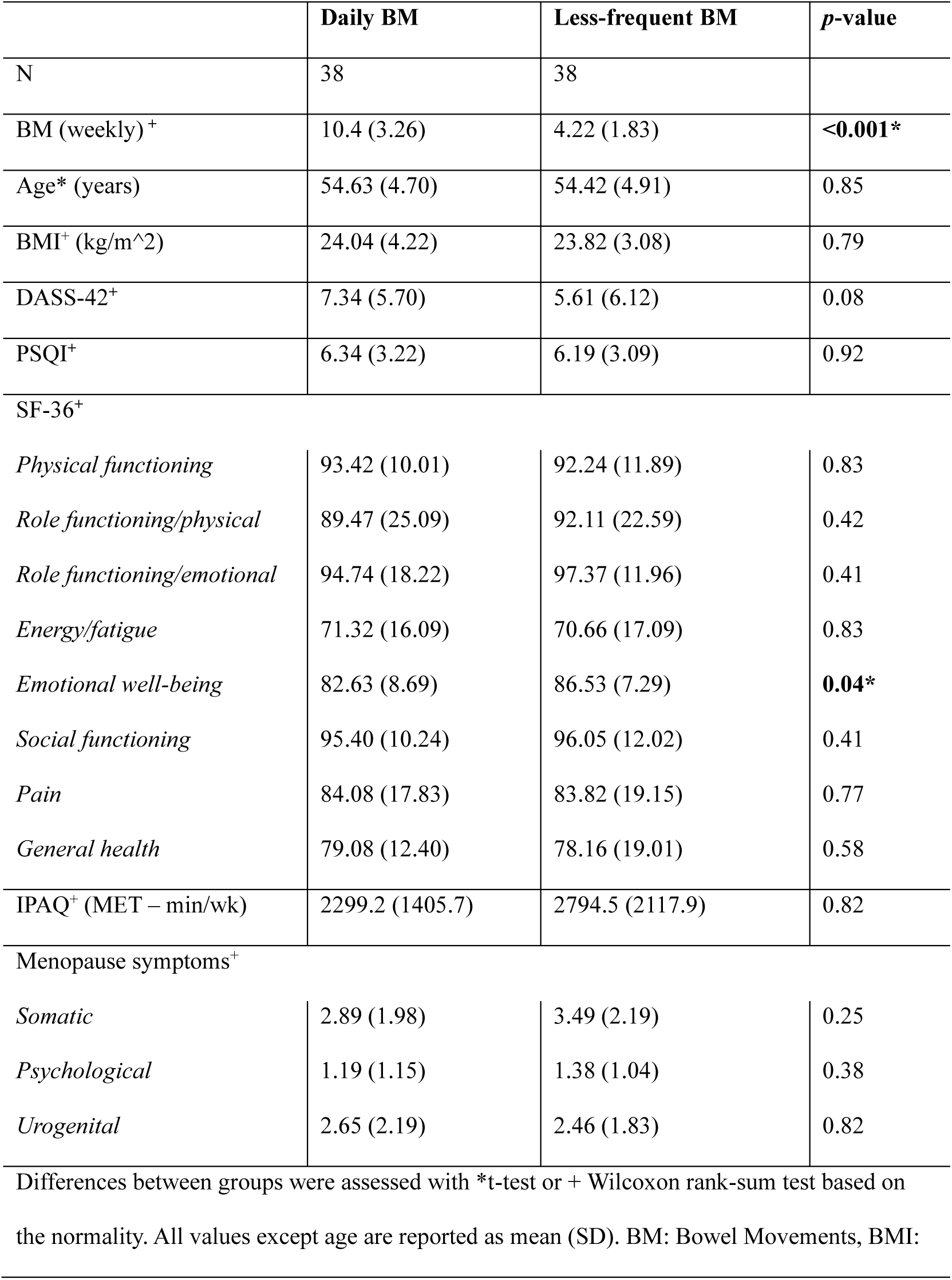
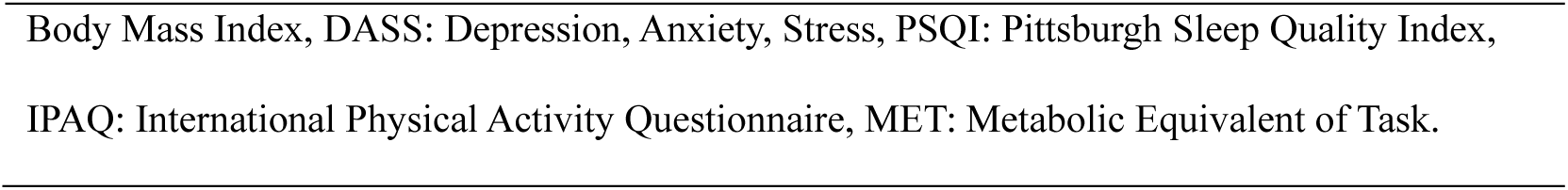
Baseline characteristics of two groups of women differing in weekly bowel movements.

There was no significant difference between groups in EEG alpha peak frequency (*p* = 0.33) and EGG gastric peak frequency (*p* = 0.93) during rest condition (**Table 2**; **Figures 3A, B**). Furthermore, the peak gastric frequency and power did not differ between the groups depicting a healthy normogastria range (∼0.05 Hz) in both groups (**Table 2**; **Figures 3C, D**). The quality control of the EGG signals confirmed 95% of the cycles fell within the normogastria range with a mean cycle duration of ∼20 seconds. The selected peak electrode had a similar spatial distribution between groups and was identified as left epigastric and infra-epigastric regions (**Figure 3C**).

**Figure 3.**
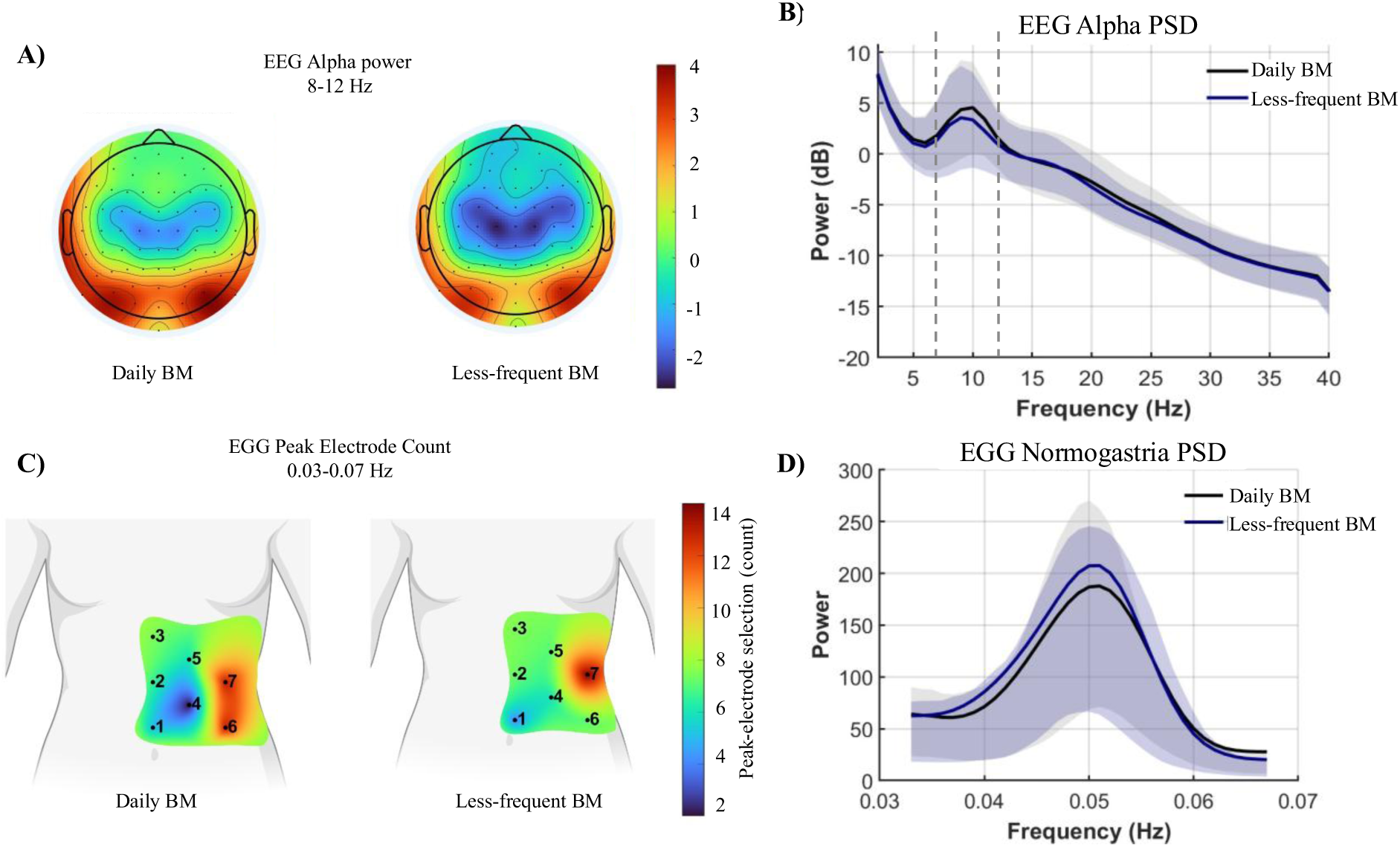
Spatial and Spectral profiles of EEG and EGG. Topographical distributions of mean EEG alpha power (A) and power-spectral density (PSD) plots display mean resting-state EEG alpha activity across the posterior region (B). EGG peak electrode count with electrode locations and electrode numbers in black (C) and PSD of abdominal peak EGG within the normogastria range (D).

**Table 2.** Baseline characteristics in Electrophysiological Measures.

|  | Daily BM | Less-frequent BM | p-value |
| --- | --- | --- | --- |
| EGG peak freq (Hz) | 0.05 (0.003) | 0.05 (0.005) | 0.93 |
| EGG gastric power (dB) | 476 (1006) | 319 (268) | 0.36 |
| EEG peak freq (Hz) | 9.74 (1.01) | 9.47 (1.31) | 0.33 |
| EEG alpha power (dB) | 7.12 (3.98) | 6.81 (4.67) | 0.75 |

Electrophysiology characteristics for both groups are reported as mean (SD). Group differences were assessed using Welch's independent samples t-test. Statistical significance was set at $p < 0.05$ .

### Women with daily bowel movements show better global cognitive performance, stronger gut-brain phase-amplitude coupling, and higher faecal SCFA levels

Notably, women in the daily BM group demonstrated lower composite-z score indicating better global cognitive performance compared to the less-frequent BM group (CI = [0.05 0.63], *p* = 0.02; **Figure 4A**). A linear regression analysis revealed that this difference remained significant after adjusting for socioeconomic status (β = 0.30, CI = [0.03 0.57], *p* = 0.03). The group difference in cognitive performance could not be ascribed to any particular cognitive domain, as no significant differences were observed in individual tests (**Table S1**).

**Figure 4.**
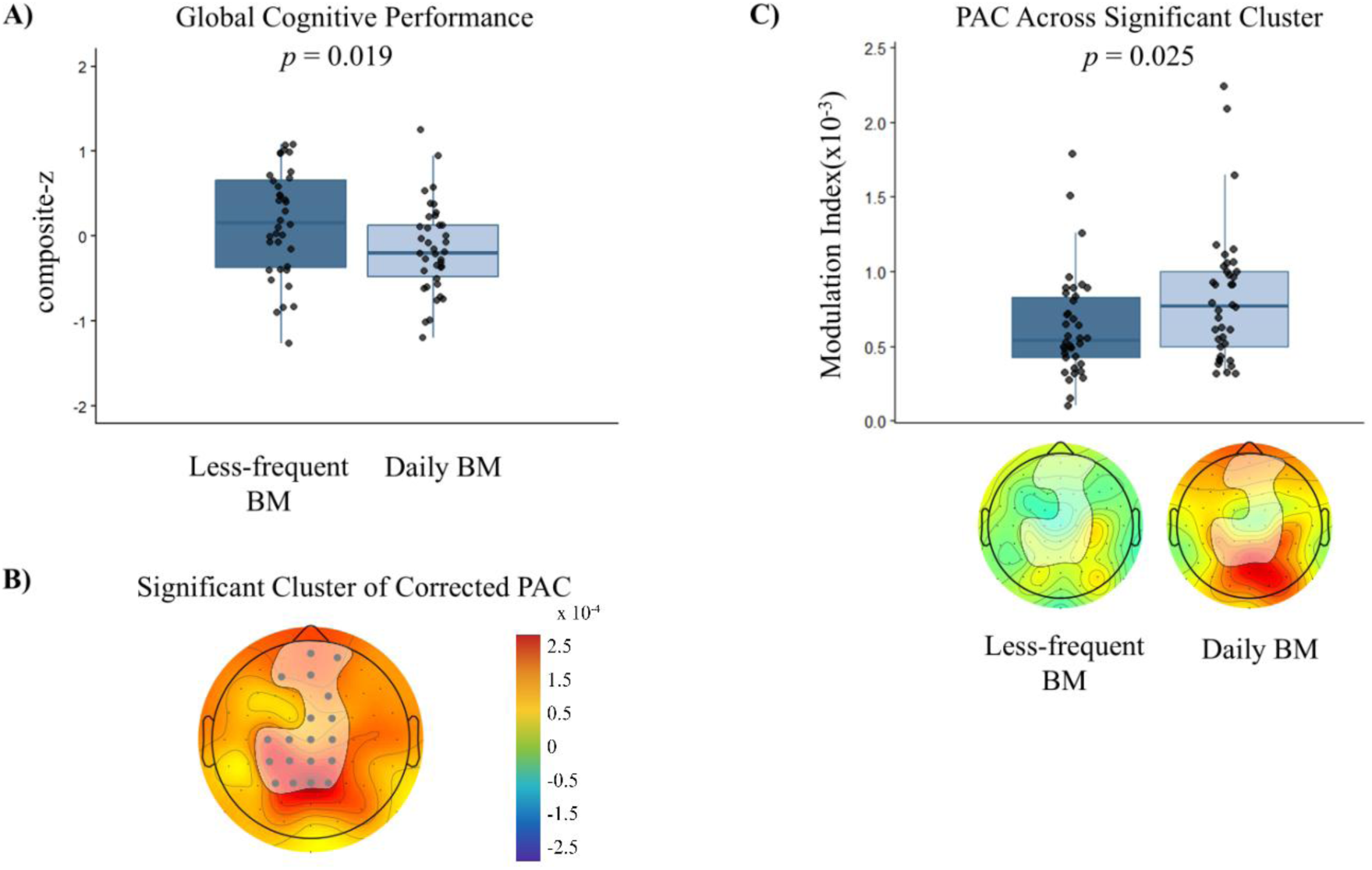
Global Cognitive Performance and Gut-brain coupling group differences. (A) Group differences in global cognitive performance. The x-axis represents the groups, and y-axis shows the cognitive composite-z score, with higher scores indicating worse performance. (B) Significant modulation index (MI) contiguous cluster identified in the analysis, overlaid in white (n = 76). (C) MI differences between daily BM (n = 38) and less-frequent BM (n = 38) groups. The x-axis represents the groups, and y-axis shows the MI. The topoplots show the average topographical distribution of MI values for each group across the significant cluster (overlaid in white). The group differences are shown in box plots, with individual participant values shown as overlaid scatter points.

Next, we investigated the phase-amplitude coupling between the phase of the gastric rhythm and brain alpha amplitude quantified by MI. Cluster-based permutation procedure indicated the presence of significant parieto-frontal clusters (ROI) (**Figure 4B**). The MI values across the ROI elucidated a significantly higher MI in the daily BM group compared to less-frequent BM group (*r* = 0.26, *p* = 0.03) (**Figure 4C**). The coupling strength also differed consistently within groups between rest and task conditions with the rest condition showing stronger coupling than the task condition (*p* < 0.005) (**Figure S4**).

Faecal SCFA profiles differed between less-frequent BM and daily BM groups. Participants with less frequent BM had significantly lower concentrations of acetate, butyrate, and propionate (**Table 3**).

**Table 3.** Faecal short-chain fatty acids (SCFA) group differences.

| | Daily BM | Less-frequent BM | $p$-value (FDR-adjusted) |
| --- | --- | --- | --- |
| Acetate (mM) <sup>+</sup> | 13.98 (7.82) | 10.32 (5.84) | 0.004 |
| Butyrate (mM) <sup>+</sup> | 3.75 (1.98) | 2.38 (1.54) | < .0005 |
| Propionate (mM) <sup>*</sup> | 3.65 (1.41) | 2.71 (1.27) | 0.004 |

SCFA levels in women with daily bowel movements and few weekly bowel movements. SCFA levels (mM) are expressed as means and standard deviation. Differences between groups were assessed with \*t-test or + Wilcoxon based on the normality. The p values are FDR corrected.

### Exploring links between gut- and brain measurements

To explore putative links between the obtained measurements, we performed correlation analyses. More frequent bowel movements were associated with better global cognitive performance (*r* = -0.29, *p* = 0.02; **Figure 5A**). Furthermore, we observed a positive correlation between BM and the MI ROI (*r* = 0.29, *p* = 0.01; **Figure 5B**). Correlation analysis further revealed positive associations between MI ROI and all three SCFAs (**Table 4**).

**Figure 5.**
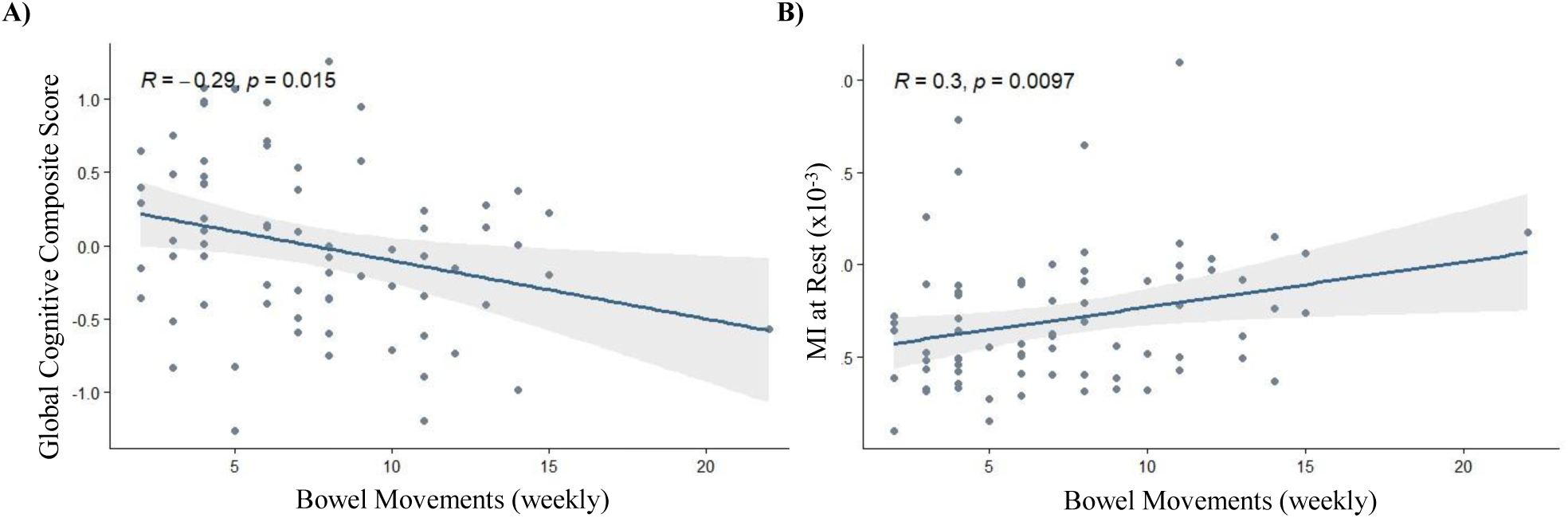
Correlation analysis of Bowel movements, Modulation Index (MI), and Cognition. Outcomes of correlating bowel movements with global cognitive performance (A) and MI at rest with bowel movements (B). Correlation plots include Spearman r and p values.

**Figure 6.**
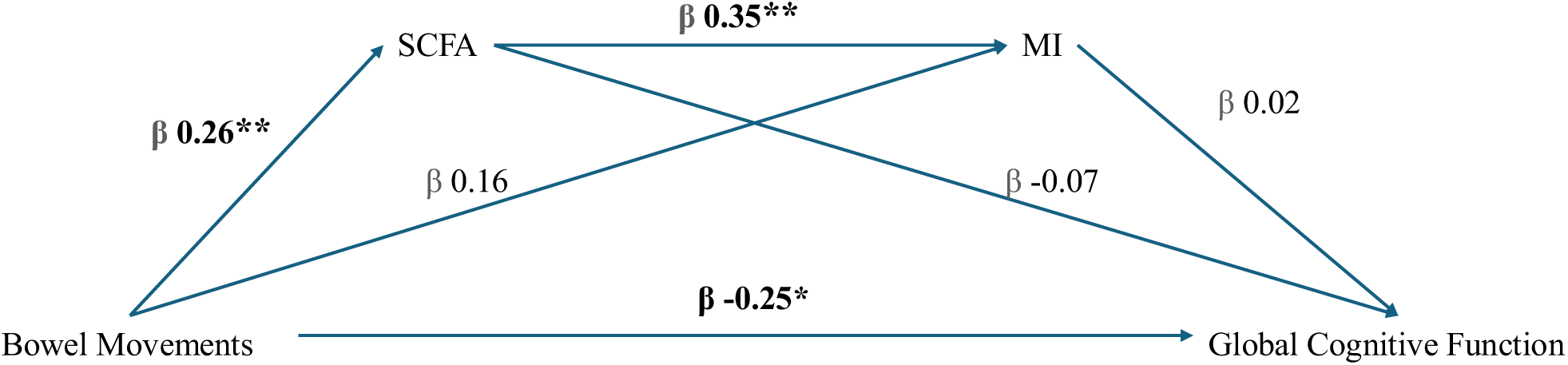
**Path model diagram**. This model shows the effect of bowel movement frequency on global cognitive performance sequentially mediated by total short chain fatty acids (SCFAs) and gut-brain modulation index (MI). Values are shown in standardized path coefficients (β). (\**p* < 0.05, \*\**p* < 0.005).

**Table 4.** Spearman correlation analysis between SCFAs and Gut-Brain Modulation Index (MI)

| | Faecal SCFA | Spearman's $r$ | Lower 95% CI | Upper 95% CI | $p$-value (FDR-adjusted) |
| --- | --- | --- | --- | --- | --- |
| Gut-Brain MI | - Acetate | 0.27 | 0.03 | 0.50 | **0.03** |
|  | - Butyrate | 0.24 | 0.01 | 0.46 | **0.04** |
|  | - Propionate | 0.32 | 0.07 | 0.52 | **0.03** |

Spearman rank correlation coefficients ( $r$ ) between modulation index and individual faecal short-chain fatty acids (SCFAs). $p$ -values were corrected for multiple comparisons using Benjamini-Hochberg false discovery rate (FDR). Significant correlations after correction are shown in bold.

### Association between Bowel Movements and Cognition via MI and SCFA

To investigate potential mediators of the association between bowel movements and cognition, we performed a mediation analysis. This analysis revealed a significant direct effect of bowel movements on cognitive performance (β = -0.25, z = -2.35, *p* = 0.02). A significant effect of bowel movements on total faecal SCFA was also observed (β = 0.26, z = 2.86, *p* < 0.005) as well as a significant effect of faecal SCFA on MI (β = 0.35, z = 3.31, *p* < 0.005). However, no significant effects of total faecal SCFA (β = -0.07, z = -0.59, *p* = 0.55) and MI (β = 0.02, z = 0.14, *p* = 0.89) on cognitive performance and bowel movements on MI (β = 0.16, z = 1.69, *p* = 0.09) were observed.

## Discussion

Here we showed that gut-brain coupling is a promising and relevant gut-brain biomarker. The test–retest reliability of MI showed a reproducible, trait-like signal with state-like fluctuations when measured at least four weeks apart. Furthermore, we found that women with daily bowel movements exhibited significantly stronger gut-brain coupling, indexed by higher parieto- frontal MI, better cognitive z-scores, and higher SCFA levels than those with less frequent bowel movements. Beyond the group differences, we observed that less frequent bowel movements were associated with lower MI values and poorer cognitive function across the whole group of participants. In addition, the faecal SCFA levels were higher in women with daily bowel movements and positively correlated with MI. The mediation path model showed that the association between bowel movements and MI was mediated by the SCFAs. However, neither MI nor SCFAs mediated the relationship between bowel movements and cognitive performance, indicating that the effect of bowel movements on cognitive performance is likely explained by other pathways.

In agreement with established literature, our gut-brain coupling results exhibit the typical spatial pattern with significant coupling in frontoparietal clusters^1,3,29,42,47^ as well as the expected reduction in coupling strength when transitioning from quiet rest to cognitive task performance^4,48^. These convergent findings underscore the robustness of gut-brain coupling as a reliable neural signature across modalities and conditions. In addition, we showed moderate stability in gut-brain coupling within participants across visits, indicating reasonable test-retest reliability. This finding contrasts with a previous study, which showed low reliability of phase-locking between the gastric rhythm and slow Blood Oxygen Level-Dependent (BOLD) fluctuations^29^. Different from MI, phase-locking value quantifies the temporal alignment of slow BOLD fluctuations and gastric rhythm. Whether EEG-EGG coupling truly is more reliable than EGG-fMRI coupling requires further investigation. There was a tendency toward higher MI values at the repeated testing session (*p* = 0.05). Although the reason for this trend remains unclear, participants may have been able to relax more during the second session, potentially increasing the MI. This phenomenon requires further investigation but suggests the importance of maintaining testing sessions as similar as possible when comparing groups. Importantly, for our baseline comparison, both groups were tested for the first time and in comparable testing surroundings such as the time of day.

The observation that bowel movements are linked to cognitive function aligns with cohort studies reporting associations between constipation and adverse cognitive outcomes in middle-aged and older adults^49,50^. To our knowledge, this is the first study to demonstrate this link in a relatively modest sample size. Critically, our use of comprehensive, domain-specific cognitive assessment and the use of objective bowel movement frequency data comparing two similar groups of women likely enhanced our ability to detect subtle cognitive associations. Importantly, gut-brain coupling strength was significantly lower in less-frequent BM group than in daily BM group, with no significant changes in non-monotonic peak frequency, gastric, or alpha rhythms.

However, as neither the MI nor faecal SCFAs mediate the effect of bowel movement on cognitive performance, other pathways, such as gut-derived hormones like peptide YY (PYY), should be investigated, as PYY primarily regulates gut motility but has also been shown to influence cognitive processes like memory, learning, and attention^51^.

Another novel aspect of our study is the integration of faecal metabolomics, revealing associations between faecal SCFAs and bowel movements, and SCFAs and gut-brain coupling strength, respectively, SCFAs are produced via gut microbiota fermenting dietary fibre and butyrate serves as the main energy source for colonic cells^52^. Fiber intake is often associated with relieving constipation, increasing bowel movement frequency, and improving stool consistency^53^. SCFAs may also increase the vagal tone through enteroendocrine cells^21^, which may explain the observed association between faecal SCFAs and gut-brain coupling strength. Even though the mediation analysis did not provide evidence that the effects of bowel movements on cognition were mediated by SCFAs, the observed associations hint towards a closely related system between the gut microbiota, colonic fermentation, bowel movements and gut-brain communication that warrants further investigation.

Several limitations should be considered, including the sample group and methods. Our study only included healthy middle-aged women, as they experience less frequent bowel movements more regularly than men or younger women^54^, but this choice limits generalizability across sexes. Our test re-test reliability analysis only included the less-frequent BM group. Given that ICC can be affected by between-subject variability, the observed moderate-reliability estimates are specific to the distribution of participants in the less-frequent BM group and may not be generalizable to daily BM group. Future studies should evaluate test-retest reliability in cohorts including daily BM groups to ensure the broader applicability of these findings. In this study, both daily BM and less-frequent BM groups had bowel habits within the relatively normal range. Extending this work to clinical populations with altered gastrointestinal function may further provide evidence regarding the directionality and causal nature of observed associations. Additionally, longitudinal interventions will be necessary to determine whether changes in gut motility and bowel movements are causally related to changes in cognition or gut-brain coupling. Such experimental modifications of gut motility and defecation frequency could either include dietary interventions or vagus nerve stimulation, which has been shown to increase gut-brain coupling^47^ and gastric motility^55^.

Another limitation of the current study is that our faecal SCFA measurements provide a limited snapshot of the microbiome function as a large fraction of SCFA is absorbed and metabolized prior to arrival in distal colon and faeces^56^. Therefore, faecal SCFA concentrations might not reliably indicate an increase in the vagal tone. However, direct measurement of vagal tone in rodent studies has shown that SCFAs, including acetate, propionate and butyrate, increase vagal afferent signalling when delivered in the small intestine, with a slower onset when compared to the other metabolites^21^. In summary, variations in bowel movement frequencies might be associated with differences in MI and cognitive performance, linking peripheral physiological rhythms to central neural dynamics. In this study we suggested a pathway model where bowel movements influence MI through SCFAs. These findings provide a basis for exploring the therapeutic and diagnostic applications of gut-brain coupling, particularly its potential to improve our understanding of the mechanisms modulating the gut-brain interactions and its role in cognitive function, mental health, and well-being

## Data & Code Availability

Deidentified participant data reported in this paper and custom scripts used in the analyses will be made publicly available at https://github.com/MovementAndNeuroscience/YourGutBrain

## Acknowledgements

We thank all the participants for their time and commitment to this study. We thank Leo Tomasevic for valuable discussions and helpful advice on electrophysiological methodology, Andreas Wulff-Abramsson for technical support, and Sofie Skov Frost for laboratory assistance with faecal samples. We also thank Trygve Tobias Granat for assistance with electrophysiological data collection.

The work was supported by Arla Food for Health and the Milk Levy Fund. The graphical abstract and Figure 1 were created using BioRender.com.

NMR data were generated though accessing research infrastructure at Aarhus University, including FOODHAY (Food and Health Open Innovation Laboratory, Danish Roadmap for Research Infrastructure).

## CRediT authorship contribution statement

**Zeynep Ertürk:** Writing – original draft, Visualization, Software, Project administration, Investigation, Data curation, Formal analysis, Validation. **Klara Nielsen:** Writing – review & editing, Methodology, Data curation, Investigation, Project administration. **Louise M.A. Jakobsen:** Writing – review & editing, Resources, Supervision. **Adam Duun Gottlieb:** Writing – review & editing, Data curation, Investigation. **Hanne Christine Bertram:** Writing – review & editing, Resources, Supervision. **Henrik M. Roager:** Conceptualization, Methodology, Supervision, Writing - review & editing, Funding acquisition. **Anke Ninija Karabanov:** Conceptualization, Methodology, Supervision, Writing – review & editing, Funding acquisition.

## Declaration of generative AI

During the preparation of this work, the authors used generative AI tools in order to improve the readability and language of the manuscript. The AI tools were not used to generate research data, perform statistical analyses, or draw scientific conclusions. The authors reviewed and edited the content as needed and take full responsibility for the content of the published article.

## Research topic(s)

Cognitive Neuroscience, Metabolomics

