## Supplementary Material for "Daily Bowel Movements are Associated with Stronger Gut-Brain Phase-Amplitude Coupling and Better Cognitive Performance"

### Appendix S1. Inclusion & exclusion criteria

**Inclusion criteria for intervention study**

Woman, 45-65 years old
Self-reported defecations every second day or less
Owns a smartphone (iOS 11.0 and later or Android 5.0 and up)
Understand Danish or English

**Inclusion criteria for the baseline sub-study**

Woman, 45-65 years old
Self-reported defecations every day
Owns a smartphone (iOS 11.0 and later or Android 5.0 and up)
Understand Danish or English

**Exclusion criteria**

Current pregnancy or lactation
Prior diagnosis of psychiatric or neurological illness
Current diagnosis of depression, anxiety, or stress
Prior diagnosis of metabolic or gastrointestinal disease (e.g., cardiovascular disease, type 1 or type 2 diabetes, chronic constipation, diarrhea, inflammatory bowel diseases (IBD) including Crohn's disease and ulcerative colitis, celiac disease, small intestinal bacterial overgrowth (SIBO), gastrointestinal obstruction, ischemic colitis, cancer, etc.)
Use of antibiotics within the last month
Use of peroral corticosteroids (inhalers excepted)
Use of medications that alter normal bowel function and metabolism (e.g., laxatives, enemas, anti-diarrheal agents, narcotics, antacids, antispasmodics, diuretics, anticonvulsants)
Use of neuroactive medications (antidepressants, anxiolytics, anticonvulsants, antipsychotics, antiparkinsonian, hypnotics, stimulants)
Concurrent participation in another trial
Any condition that makes the project responsible researcher doubt the feasibility of the volunteer´s participation

#### Appendix S2. Defecation diary

### Defecation diary - before visit 1

*Must be completed for all stools in the week before visit 1.*

| **Date** (day-month-year) | **Time** (hour:minute) | **Bristol scale** (type, number) | **Presence of sweet corn in stool** (yes/no) | **Stool collected** (tick if yes**)** |
|---|---|---|---|---|

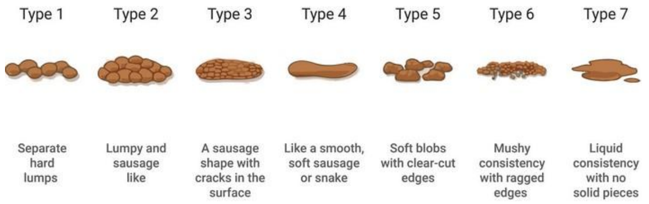

##### Registration of sweet corn intake

*Sweet corn is consumed four days before the visit. Remember to note if sweet corn is observed in the stool in the following days.*

| | **Date** (day-month-year) | **Time** (hour:minute) |
|---|---|---|
| **100 g sweet corn (eaten before dinner four days before the visit)** | | |

*The first stool after eating sweet corn must be collected in the provided box, stored in your own freezer, and brought to the institute using the supplied cooling bag and freezing elements.*

#### Appendix S3 - Questionnaire regarding menopause and lifestyle

**Questions regarding menopause**

a. Are you or have you been in menopause? _____ Yes _____ No _____ Don’t know

b. If you answered ‘Yes’ or ‘Don’t know’ to question a. please note when your last period ended Date __________

c. Please fill out the questionnaire regarding menopause symptoms below.

**Questions regarding lifestyle**

d. How many drinks containing alcohol do you drink per week __________ Alcoholic drinks/week

e. Do you smoke?
_____ Yes _____ No

f. If you answered yes to question e. Please note how many cigarettes you smoke per day. __________ Cigarettes/day

Please note how much the following symptoms bother you by marking an X in the box you find most fitting. If you mark 2 or 3 for some of the symptoms, you may need to speak to your doctor.

| **SYMPTOM** | **Not at all** | **A little** | **Quite a bit** | **All the time** | **Comments** |
|---|---|---|---|---|---|
| | 0 | 1 | 2 | 3 | |
| **Hot Flushes** – Your body feels hot and sweaty during the day or night. | | | | | |
| **Cold Flushes** – Your body feels very cold during the day or night | | | | | |
| **Heart Palpitations** – heartbeat is different to normal and you can feel anxious or worried | | | | | |
| **Irritability** – Things and people make you angry or annoyed | | | | | |
| **Trouble Sleeping** – your normal sleep routine is not as it was | | | | | |
| **Irregular periods or stopped periods** -<br>Your period is stopped, or you sometimes have your period but also experience missing a month or two. * | | | | | |
| **Low Sex Drive** – you do not want to have sex as much as you used to or might not want to have sex at all | | | | | |
| **Poor Concentration** – find it hard to concentrate like you used to | | | | | |
| **Incontinence** – sometimes not getting to the toilet quick enough | | | | | |
| **Sore Breasts** | | | | | |
| **More facial Hair** – random hairs may appear on your chin; they seem to grow overnight | | | | | |
| **Dizziness** – you experience dizziness when you have not normally experienced this | | | | | |
| **Changed body smell** – you smell different to normal | | | | | |
| **Osteoporosis –** weakened bones, tends to result in fractures | | | | | |
| **Vaginal dryness or tearing of the skin around the vagina (also known as Atrophy)**. Symptoms can include itching and can lead to bleeding when wiping after a visit to toilet and painful Sex. * | | | | | |

*Wording changed from original questionnaire.

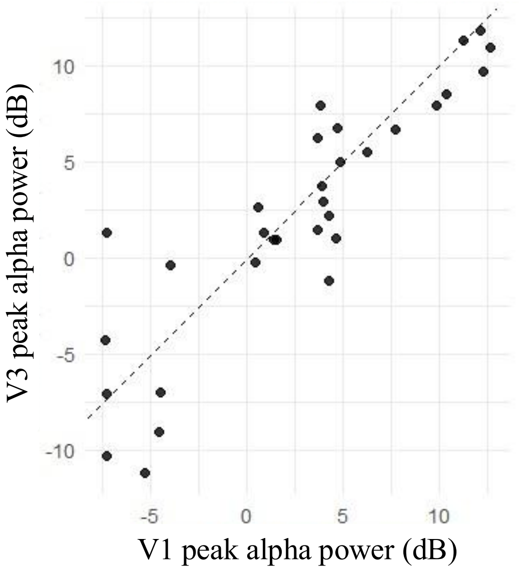

**Figure S1. EEG alpha test-retest reliability.** Scatter plot of EEG alpha power values measured at baseline (V1; a-xis) and retest (V3; y-axis).

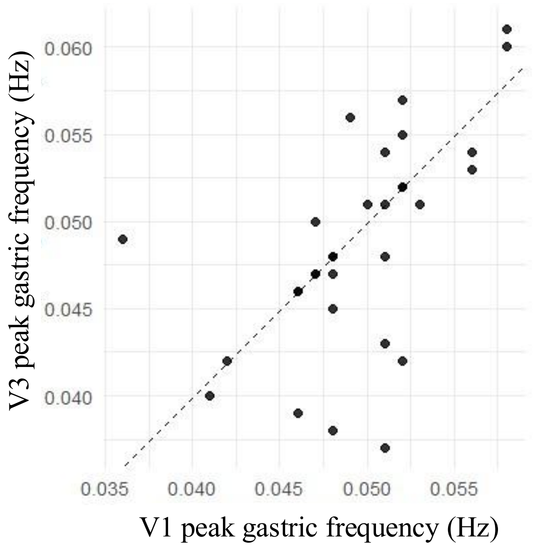

**Figure S2. EGG gastric frequency test re-test reliability.** Scatter plot of EGG peak gastric frequency values measured at baseline (V1; a-xis) and retest (V3; y-axis).

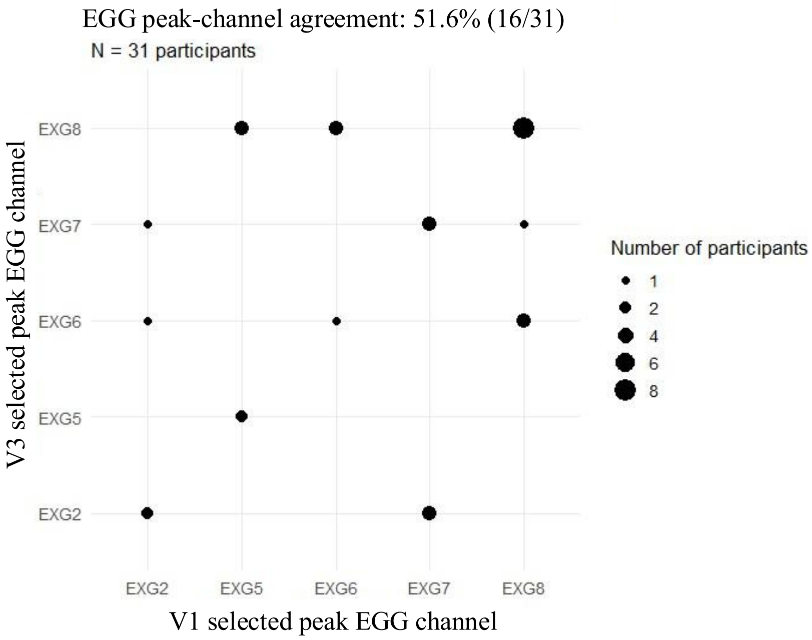

**Figure S3. Test-retest agreement of the selected EGG maximum channel.** Maximum EGG channel at baseline (V1) I splotted against the maximum EGG channel at restest (V3). Point size represents the number of participants with each V1-V3 channel combination. Points on the diagonal indicate exact agreement in the selected maximum channel between visits.

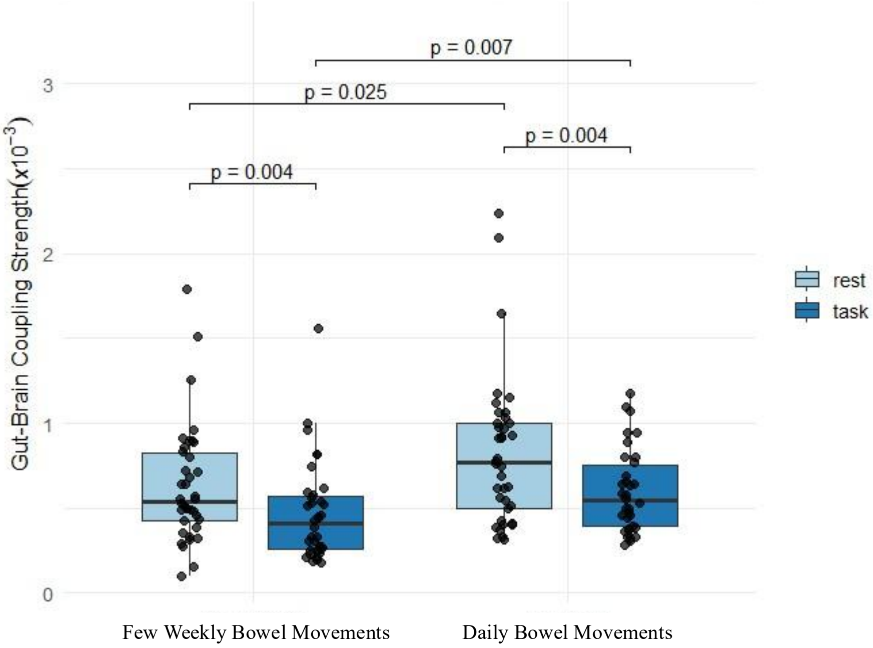

**Figure S4. Modulation Index Task versus Rest.** Boxplots show the distribution of phase-amplitude coupling modulation indes (MI) values, with individual participant observations overlaid. Rest and task conditions were compared using a Wilcoxon signed-rank test in the daily-BM group and a paired t-test in the less-frequent BM group. Between- group differences were assessed using Wilcoxon rank-sum tests for both rest and task conditions. Corresponding p-values are also shown above the comparisons.

**Table S1. CANTAB Group Differences Selected Measures**.

| Selected Measure | Reference (SD) | Low BM (SD) | p_value |
|---|---|---|---|
| N | 38 | 36 | |
| MOTML$^{+}$ | 595 (87) | 634 (122) | 0.37 |
| PALTEA$^{+}$ | 10.6 (9.78) | 14.3 (10.4) | 0.05 |
| PALTEA4$^{+}$ | 0.55 (1.01) | 0.69 (1.49) | 0.98 |
| PALTEA6$^{+}$ | 3.24 (3.68) | 3.64 (3.68) | 0.39 |
| PALTEA8$^{+}$ | 6.66 (6.46) | 9.92 (7.49) | 0.05 |
| RVPPFA$^{+}$ | 0.00 (0.00) | 0.01 (0.01) | 0.14 |
| RVPTFA$^{+}$ | 1.58 (1.91) | 2.42 (2.47) | 0.14 |
| EBTBP* | 8.85 (1.46) | 9.00 (1.33) | 0.65 |

MOTML: reaction time in motor screening, PALTEA: Adjusted errors in paired associates learning task to assess visual memoty and learning including 4$^{th}$, 6$^{th}$ and 8$^{th}$ stage indicating increasing difficulty with increased number of shapes to remember, RVPPFA: Probability of false alarms in rapid visual information processing. task to assess sustained attention, RVPTFA: Total false alarms in rapid visual information processing task, EBTBP: bias point in the emotional bias task to assess perceptual bias in facial emotion perception (this measure is not included in the composite-z score). *t-test or + Wilcoxon based on the normality.
